# Distinct Mechanisms Underlie Electrical Coupling Resonance and Its Interaction with Membrane Potential Resonance

**DOI:** 10.1101/2023.01.11.523652

**Authors:** Xinping Li, Omar Itani, Dirk M. Bucher, Horacio G. Rotstein, Farzan Nadim

**Affiliations:** Department of Biological Sciences, New Jersey Institute of Technology, Newark, NJ, USA

**Keywords:** Oscillation, Central Pattern Generator, Stomatogastric, Gap Junctions

## Abstract

Neurons in oscillatory networks often exhibit membrane potential resonance, a peak impedance at a non-zero input frequency. In electrically coupled oscillatory networks, the coupling coefficient (the ratio of post- and prejunctional voltage responses) could also show resonance. Such coupling resonance may emerge from the interaction between the coupling current and resonance properties of the coupled neurons, but this relationship has not been clearly described. Additionally, it is unknown if the gap-junction mediated electrical coupling conductance may have frequency dependence. We examined these questions by recording a pair of electrically coupled neurons in the oscillatory pyloric network of the crab *Cancer borealis*. We performed dual current- and voltage-clamp recordings and quantified the frequency preference of the coupled neurons, the coupling coefficient, the electrical conductance, and the postjunctional neuronal response. We found that all components exhibit frequency selectivity, but with distinct preferred frequencies. Mathematical and computational analysis showed that membrane potential resonance of the postjunctional neuron was sufficient to give rise to resonance properties of the coupling coefficient, but not the coupling conductance. A distinct coupling conductance resonance frequency therefore emerges either from other circuit components or from the gating properties of the gap junctions. Finally, to explore the functional effect of the resonance of the coupling conductance, we examined its role in synchronizing neuronal the activities of electrically coupled bursting model neurons. Together, our findings elucidate factors that produce electrical coupling resonance and the function of this resonance in oscillatory networks.

## 1 Introduction

In oscillatory circuits, neurons and synapses are subject to inputs that often span a range of frequencies. Whether they respond more favorably in one frequency range, and whether such frequency selectivity can be altered in different states, may impact the dynamics of the circuit output. Many neurons exhibit a frequency-dependent property known as membrane potential resonance, characterized as a maximal subthreshold impedance at a non-zero (resonance) frequency (Hutcheon and Yarom, 2000). When measured with oscillatory current injection, this corresponds to the voltage amplitude response being maximal to oscillatory current input at that frequency. Membrane potential resonance typically arises through interactions of passive properties of the neuron and the kinetics of voltage gated ionic currents (Hutcheon and Yarom, 2000). The resonance frequency of neurons has been shown to correlate with the network frequency in several systems (Wu et al., 2001; Bykhovskaia et al., 2004; Tohidi and Nadim, 2009; Moca et al., 2012). Membrane potential resonance is one form of preferred frequency response observed in neural circuits, but other circuit properties such as synaptic strengths and firing rate can also have a preferred frequency at which the output is maximized and such preferred frequencies are also often termed resonance (Izhikevich et al., 2003; Richardson et al., 2003; Drover et al., 2007; Ledoux and Brunel, 2011; Tseng et al., 2014; Rau et al., 2015; Stark et al., 2022).

In neural circuits coupled through gap junction-mediated electrical coupling, any input that causes membrane potential oscillations in one neuron could produce oscillations in its coupled partners (Landisman et al., 2002; Long et al., 2004). In electrically coupled networks where individual neurons exhibit membrane potential resonance, both the postjunctional neuron’s membrane potential and the coupling coefficient (the ratio of post- and prejunctional voltages) can also exhibit preferred frequency responses (Curti et al., 2012; Stagkourakis et al., 2018). However, it is not known if coupling resonance reflects the properties of the electrical coupling, those of the coupled neurons, or if it emerges from the interaction between the two. Electrical coupling is an important factor in generating neural oscillations (Posłuszny, 2014; Coulon and Landisman, 2017; Traub et al., 2018; Alcamí and Pereda, 2019) and, as we showed in a previous study, membrane potential resonance can directly influence the network oscillation frequency through electrical coupling (Chen et al., 2016). It is therefore important to understand how resonance properties of neurons can interact through electrical coupling.

We examined this question by recording pairs of electrically coupled neurons that show resonance in the oscillatory pyloric network of the crab, *Cancer borealis*. This circuit includes two bursting pyloric dilator (PD) neurons that are known to exhibit membrane potential resonance at a frequency close to the pyloric circuit oscillation frequency (Tohidi and Nadim, 2009; Fox et al., 2017). These two neurons are strongly electrically coupled to each other and, during normal activity, exhibit synchronous slow-wave oscillations that support their bursting activity (Marder and Eisen, 1984). We took advantage of the fact that we could examine the PD neurons’ membrane potential resonance and their coupling properties simultaneously to quantify the frequency dependent properties of the neurons, the coupling coefficient, and the coupling current (measured in voltage clamp). We found that all three components exhibit frequency selectivity, but with distinct preferred frequencies. Although resonance in the coupling coefficient has been previously reported, this is, to our knowledge, the first report of resonance in the coupling current.

We used mathematical analysis and computational modeling to explain the mechanism underlying resonance in the coupling coefficient, and what factors determine its resonance frequency. We then examined potential circuit mechanisms that may give rise to resonance in the coupling current and explored how such a resonance may influence network synchronization.

## 2 Materials and Methods

### 2.0 Preparation and Electrophysiology Recordings

All experiments were performed on wild-caught adult male crabs (*Cancer borealis*) purchased from local seafood suppliers in Newark, NJ. Prior to experiments, animals were kept in artificial sea water tanks at 13 °C. Before dissection, crabs were anesthetized by placing them on ice for 30 min. The STNS was dissected out following standard protocols (Blitz et al., 2004; Tohidi and Nadim, 2009), placed in a Petri dish coated with clear silicone elastomer (Sylgard 184; Dow Corning) and superfused with *C. borealis* saline, containing (in mM) 11 KCl, 440 NaCl, 13 CaCl_2_, 26 MgCl_2_, 11.2 Trizma base, and 5.1 maleic acid (pH =7.4 –7.5). A petroleum jelly well was built around the STG for constant superfusion of chilled (10-12 °C) saline during the experiment.

PD neurons were identified by their characteristic intracellular waveforms and by matching their activities to the spikes on the corresponding motor nerves. Extracellular activities of motor nerves were recorded with a differential AC amplifier (Model 1700; A-M Systems), using stainless-steel pin wire electrodes placed inside and outside of small petroleum jelly wells built around the nerves. Intracellular recordings, current clamp and voltage clamp experiments were done with Axoclamp 900A amplifiers (Molecular Devices). The STG was desheathed and the neuron cell bodies were impaled with sharp glass electrodes, prepared with a Flaming-Brown P-97 Puller (Sutter Instruments) and filled with 0.6 M K_2_SO_4_ + 20 mM KCl solution (15-30 MΩ electrode resistance). All electrophysiological data were digitized at 5-10 KHz with a Digidata 1440A data acquisition board (Molecular Devices).

### 2.1 Measuring Electrical Coupling Resonance and Membrane Potential Resonance

We measured the membrane potential and electrical coupling resonance in pairs of PD neurons, in both current clamp experiments and voltage clamp experiments, with dual two-electrode recordings. In all experiments, we recorded the voltage in both the pre- and the postjunctional neurons (*V*_*pre*_ and *V*_*post*_) and the current injected into them (*I*_*pre*_ and *I*_*post*_). In current clamp experiments, a ZAP (Impedance Amplitude Profile) current was injected into the prejunctional neuron and produced oscillation in both *V*_*pre*_ and *V*_*post*_. The ZAP function was given by

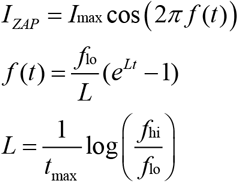

where *f*(*t*) swept a range of frequencies as a function of time, *t*, from *f*_lo_ = 0.1 Hz to *f*_hi_ = 4 Hz. *I*_max_ = 3nA and produced a *V*_*pre*_ roughly ranging from –60 mV to –30 mV. *T* is the total duration of the ZAP waveform which, in most trials was at least 100 s. Additionally, to avoid transients, we always started the ZAP function with 2 pre-cycles of a sinusoidal current applied at the lowest frequency (*f*_lo_ = 0.1 Hz) that smoothly transitioned into the ZAP waveform. When measuring in voltage clamp, the same ZAP function was applied to the prejunctional voltage *V*_*pre*_ to force it to alternate between –60 and –30 mV, while the postjunctional neuron was held at a constant voltage of *V*_*post*_ = –60 mV. The prejunctional impedance (*Z*_*pre*_), the postjunctional impedance (*Z*_*post*_), the coupling coefficient (*CC*) and the coupling conductance (*G*_*c*_) were calculated as shown in Table 1.

**Table 1.**
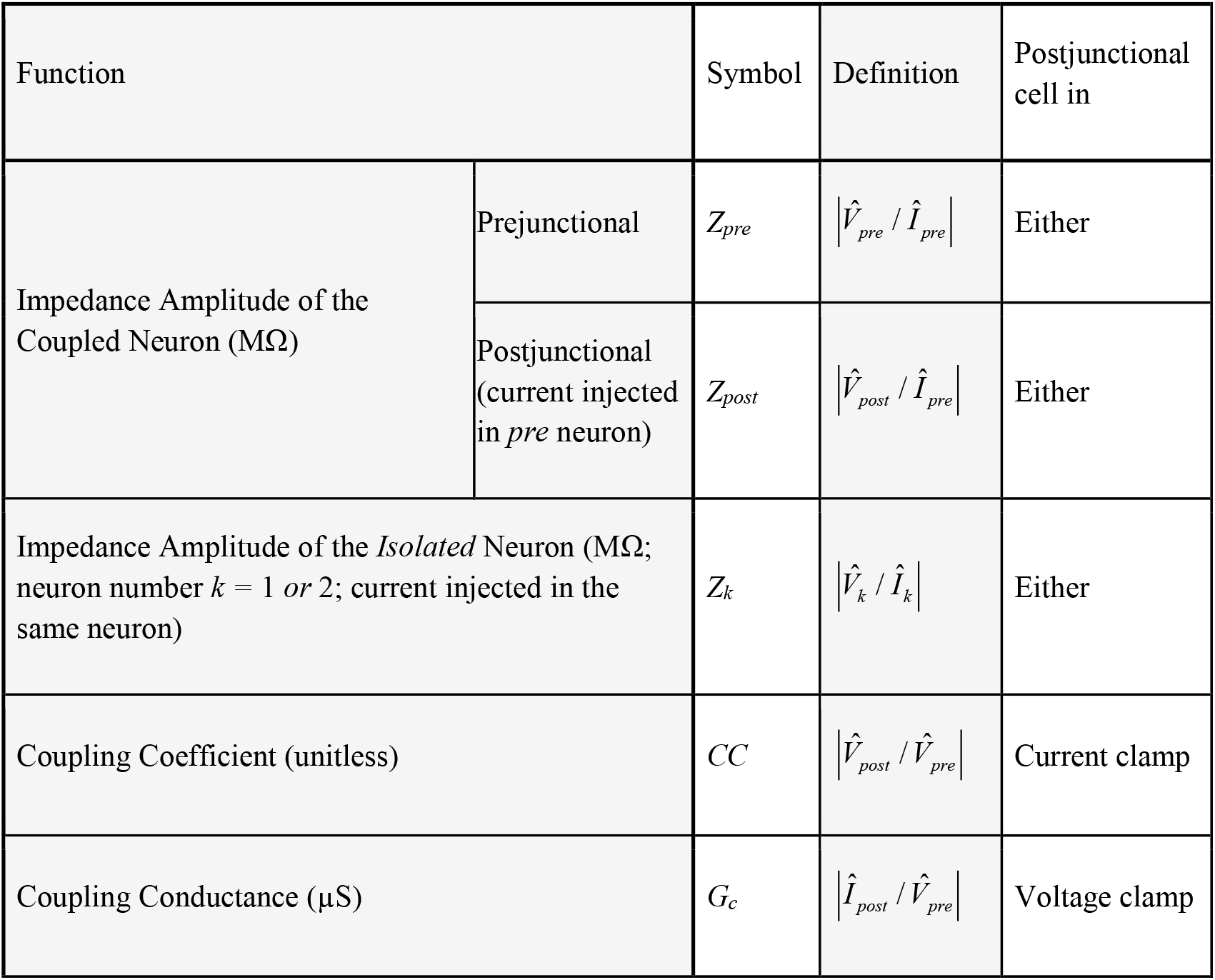
List of notations. All symbols in the table are functions of the input frequency *f*. The symbol 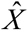 refers to the Fourier transform of *X*. In this manuscript we use the symbols below to denote the norm (|·|) of the complex values obtained by the Fourier transforms.

All factors measured as a function of frequency, in current or voltage clamp, were fit with a sixth-degree polynomial in MATLAB (MathWorks) and the resonance frequency and amplitude were estimated as the peak amplitude of the fit curve and the frequency at which the maximum amplitude was achieved.

All experimental measurements involving electrical coupling were done in the presence of 100 nM tetrodotoxin citrate (TTX; Biotium) saline to block action potentials as well as the descending neuromodulatory inputs, and 5 µM picrotoxin (PTX; Sigma) to block chemical synapses within the STG, all of which are inhibitory.

### 2.2 Data and Statistical Analysis

All experimental data analysis was done using scripts written in MATLAB, and statistical comparisons were done in SigmaPlot 12 (SyStat Software Inc.). Critical significance level was set to α = 0.05. Unless otherwise indicated, all error bars in the figures represent standard error of the mean.

### 2.3 Model of coupled resonant neurons

We made biophysical models of coupled resonant neurons of Fig. 5, using single compartment neurons having the Hodgkin-Huxley type currents given in Table 2. The model structure and parameters for the model neurons were implemented from the PD neuron resonance properties as previously described (Fox et al., 2017). All simulations were performed in NEURON 8.0 through the Python 3.8 interface. Analyses were conducted through custom Python scripts using scipy 1.5 and numpy 1.19 packages. All simulations for this study are available on https://github.com/fnadim/ECouplingResonance.

**Table 2.**
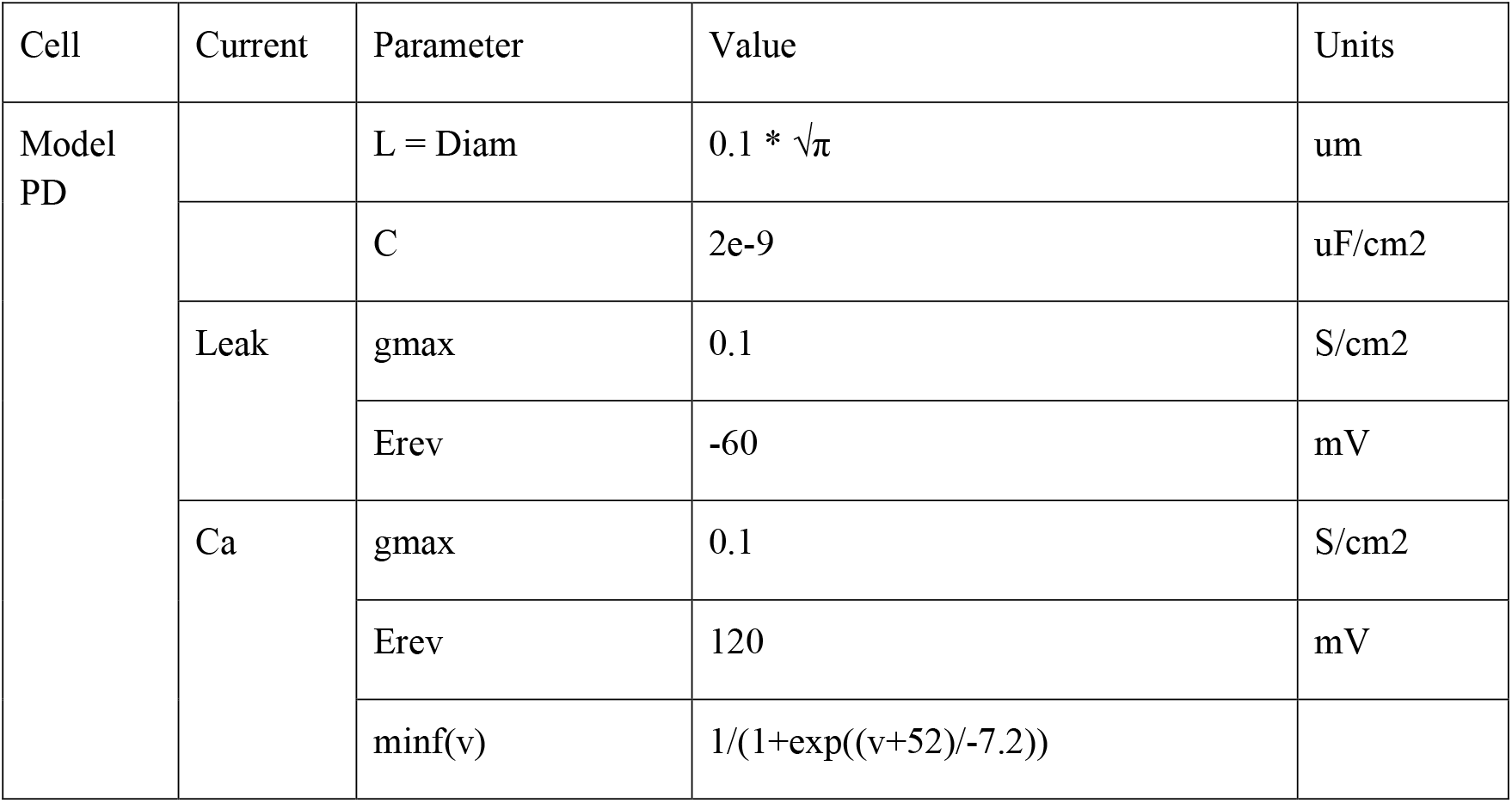
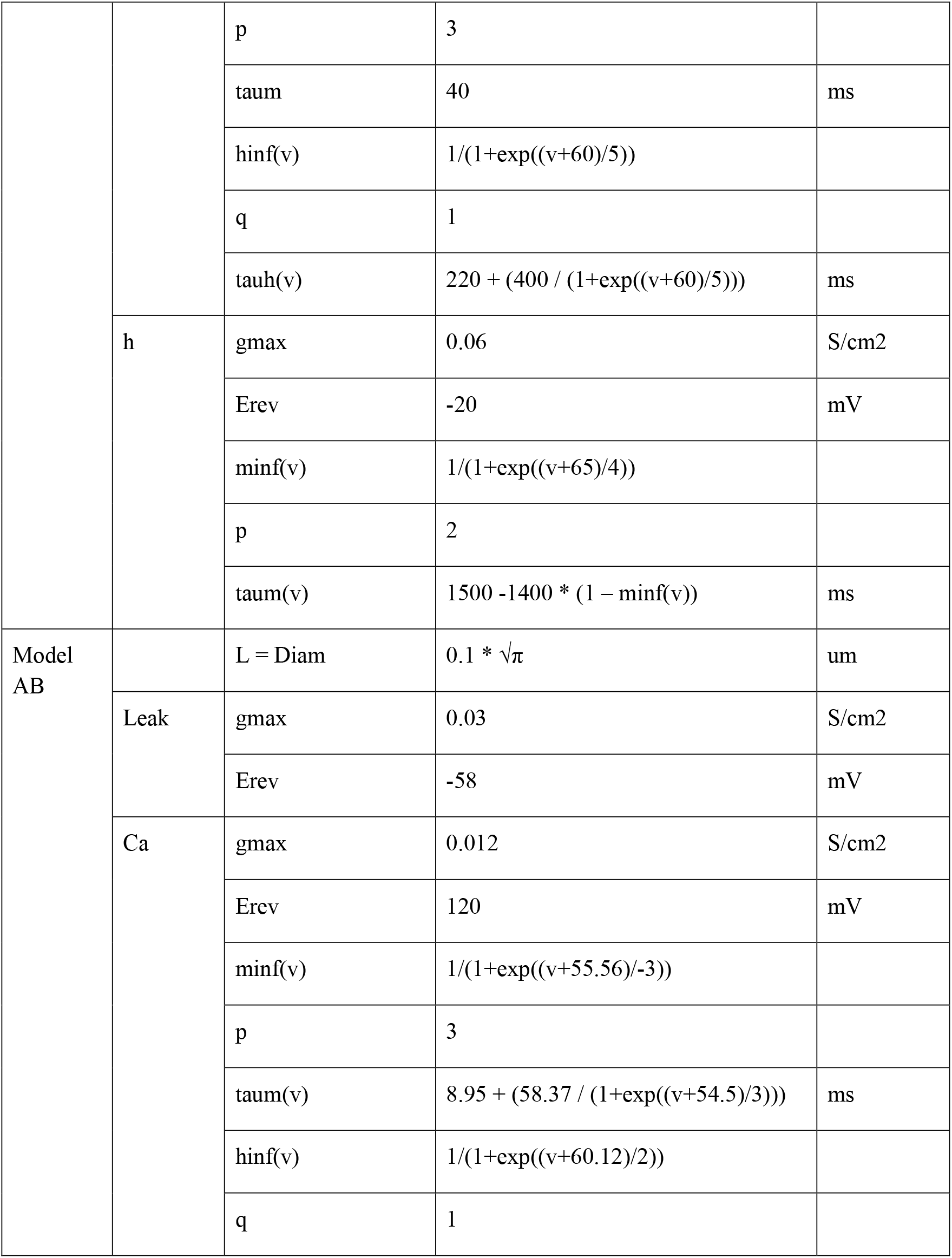
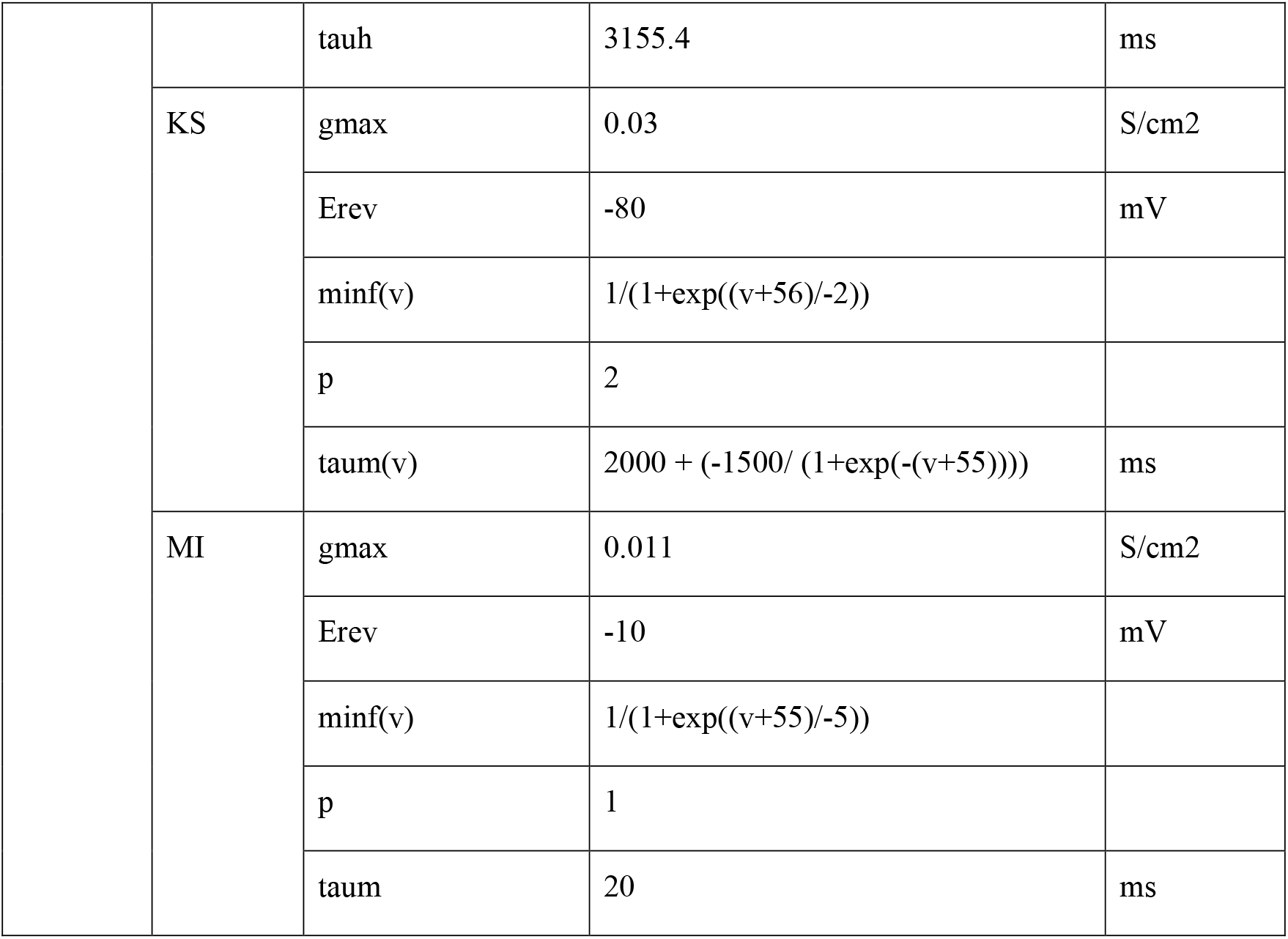
Parameters of the coupled resonant neurons.

### 2.4 Model of coupled bursting neurons

The model consisted of two neurons coupled with symmetric electrical coupling. Each neuron was built as a two-compartment biophysical model, consisting of a soma/neurite (*SN*) and an axon (*A*) compartment. The soma/neurite compartment included a leak and a low-threshold (T-type) inactivating calcium current, which effectively made it a calcium spike oscillator (Torben-Nielsen et al., 2012). The axon compartment included Hodgkin-Huxley type leak, fast sodium and delayed rectifier potassium currents, which allowed it to spike but only when the input from the soma/neurite compartment produced a calcium spike. The combination produced a bursting neuron. The neuron obeyed the following standard Hodgkin-Huxley type current balance equations:

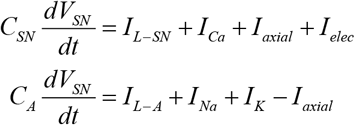

where *C*_*x*_ and *I*_*L*−*x*_ = *g*_*L*−*x*_ (*V* − *E*_*L*−*x*_) denote the membrane capacitance and leak current of the compartments (*x = SN* or *A*), *I*_*axial*_ = *g*_*axial*_ (*V*_*SN*_−*V*_*A*_) and *I*_*axial*_ = *g*_*elec*_ (*V*_*SN*_−*V*_*SN* 2_) where *V*_*SN2*_ is the voltage of the other neuron’s *SN* compartment. The ionic currents are given as

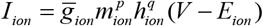

where *ion* = *Ca, Na* or *K*, 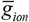 is the maximal conductance, and *m*_*ion*_ and *h*_*ion*_ denote the activation and inactivation gating variables governed by

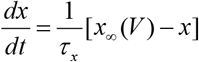

(*x* = *m*_*ion*_ or *h*_*ion*_). The activation and inactivation powers, *p* and *q*, are nonzero integers. The model equations and parameters are provided in Table 3. The parameters of the two neurons were chosen so that, in isolation, their bursting frequencies differed by about 10%.

**Table 3.**
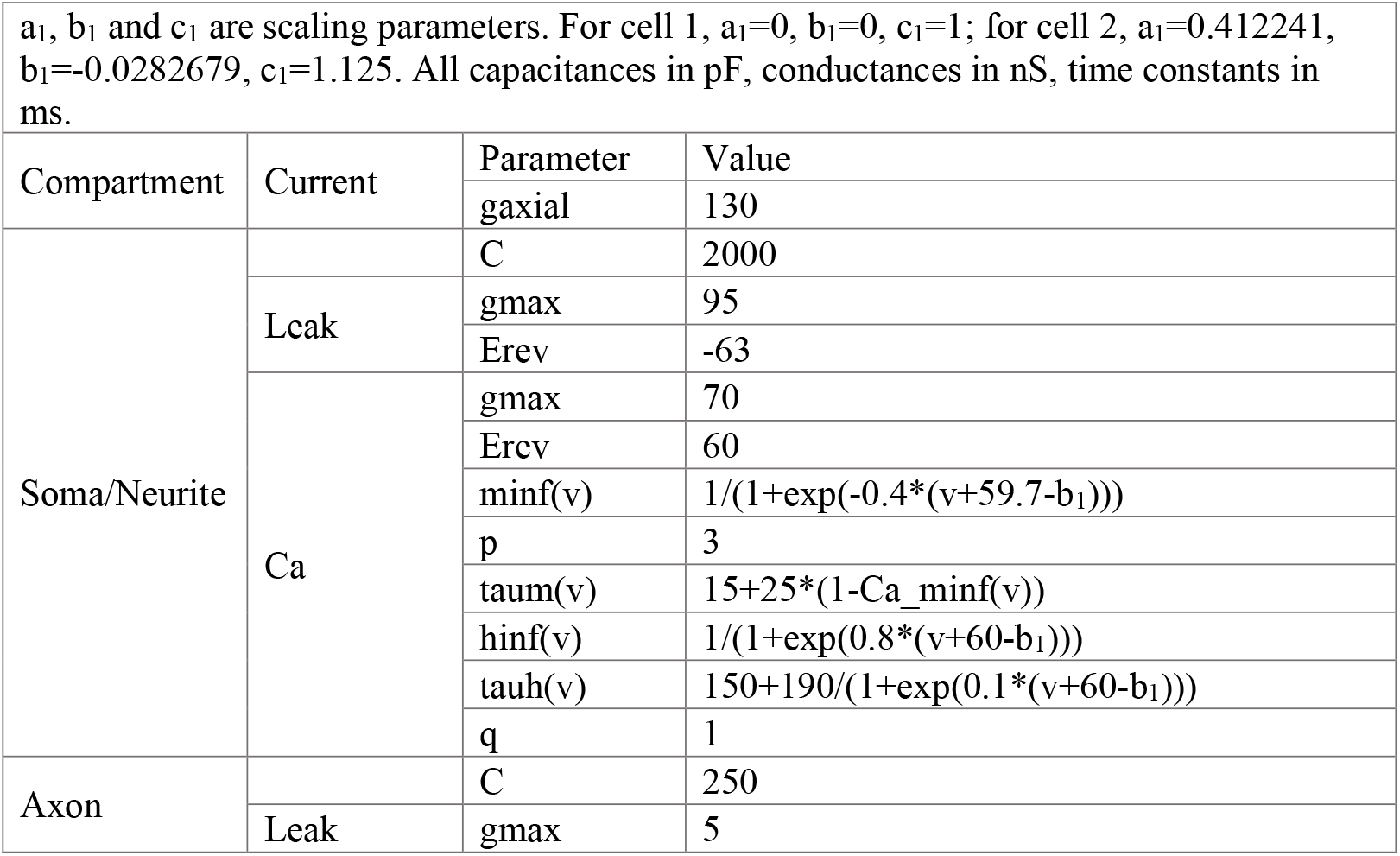
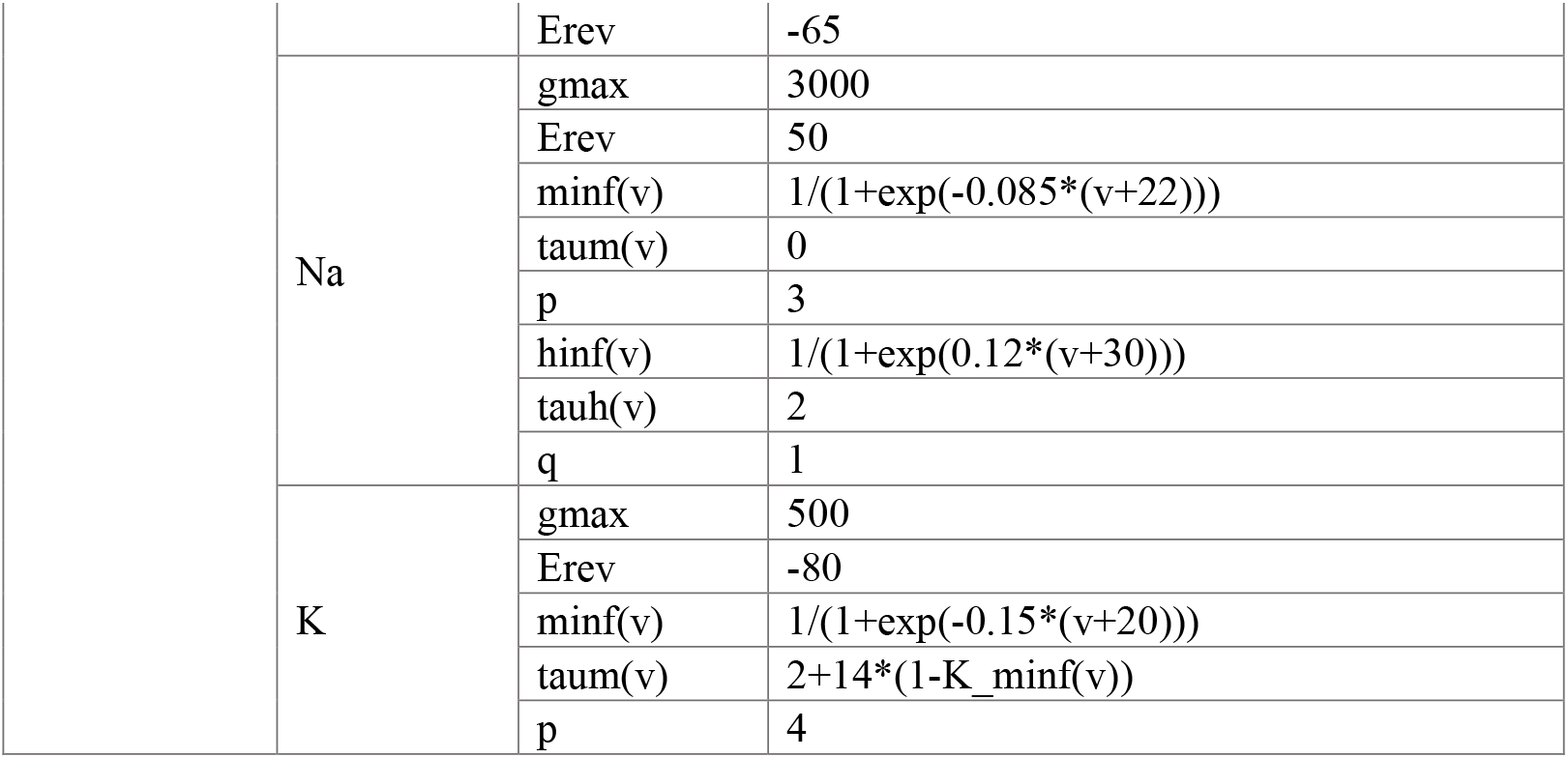
Parameters of the coupled bursting neurons.

The *G*_*c*_ frequency profile was modeled to show resonance at *f* = 0.75 Hz according to the following equation:

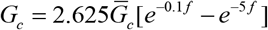

where 2.625 is a scaling factor so that 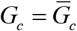 at the resonance frequency.

Simulations were done in C, using a 4^th^ order Runge-Kutta numerical integrator. The two cells always started with identical initial conditions and each run was 25 s. A 15 s window, ending 1 s before the simulation end (to remove filtering artifacts), was used for measurements of synchrony. The two voltage waveforms were sampled at 1 KHz The Slow waveform was obtained by low-pass filtering the waveforms with a moving average window of length 81 ms. The Fast waveform was obtained as the difference between the Full waveform and the Slow waveform. The level of synchrony was measured as, R^2^, the square of the correlation coefficient between the (Full, Slow or Fast) waveforms of the two cells in this time window. All analysis was done in MATLAB (MathWorks).

## 3 Results

### 3.0 The coupling coefficient between the PD neurons exhibits resonance at a distinct frequency from their membrane potential resonance

The two PD neurons are very similar in their ionic current expression and anatomical structure and therefore considered to be functionally equivalent, if not identical (Marder and Eisen, 1984; Bucher et al., 2005; Schulz et al., 2006). During normal pyloric activity, these two neurons exhibit synchronous slow-wave oscillations that support their bursting activity (Fig. 1A). This synchronous activity arises primarily from their electrical coupling to one another and to the pyloric pacemaker, the anterior burster (AB) neuron (Marder and Eisen, 1984). The electrical coupling strength between the two PD neurons can be determined in the classical way as the coupling coefficient (*CC*), measured as the ratio of the voltage change of the postjunctional neuron to that of the prejunctional neuron (Fig. 1B):

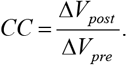

**Figure 1.**
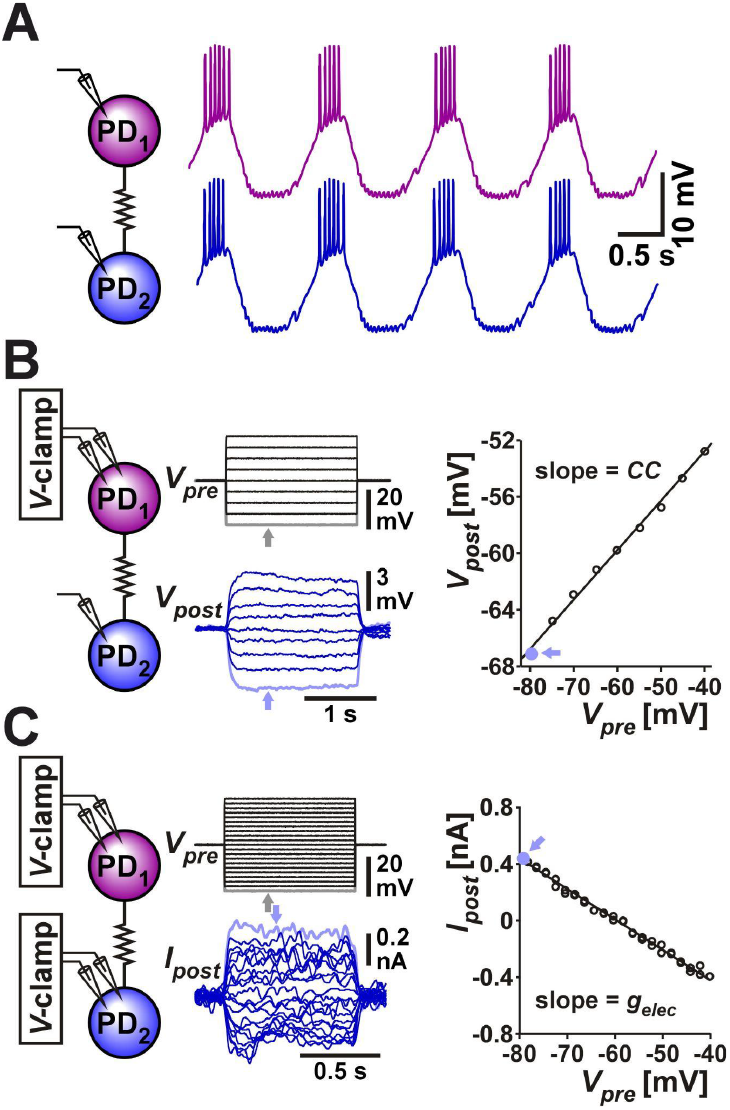
The two PD neurons produce synchronized slow wave bursting due to their strong electrical coupling. **(A)** Somatic recording of the two PD neurons shows that they produce bursting oscillations that are synchronized in their slow-wave activity. **(B)** Measurement of coupling coefficient between the two PD neurons. The prejunctional PD_1_ neuron is voltage clamped with steps ranging from -80 to -40 mV from a holding potential of -60 mV. The postjunctional PD_2_ neuron membrane potential is recorded in current clamp. The coupling coefficient *CC* is measured as the slope of the linear fit to the values of *V*_*post*_ plotted vs. *V*_*pre*_. Each data point is the mean value of voltage during the step, as seen in the grey point, corresponding to the lowest steps (arrows). **(C)** Measurement of coupling conductance between the two PD neurons. The prejunctional PD_1_ neuron is voltage clamped as in panel B, while the postjunctional PD_2_ neuron is voltage clamped at a steady holding potential of - 60 mV (not shown). The coupling conductance *G*_*c*_ is measured as the slope of the linear fit to the values of *I*_*post*_ plotted vs. *V*_*pre*_. Each data point is the mean value the step, as seen in the grey point, corresponding to the lowest *V*_*pre*_ and highest *I*_*post*_ steps (arrows).

A more direct measure of the strength of coupling, which does not depend on the input resistance of the postjunctional neuron can be obtained by voltage clamping both neurons, stepping the voltages of the (arbitrarily-designated) prejunctional neuron and measuring the current flow to the postjunctional cell. The coupling conductance (*G*_*c*_) can be measured as (Fig. 1C):

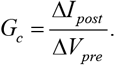

The PD neurons are bursting oscillators and, additionally, these neurons show membrane potential resonance at a frequency correlated with their burst frequency (Tohidi and Nadim, 2009; Tseng and Nadim, 2010; Fox et al., 2017). We were interested in knowing whether the coupling strength between the two PD neurons (the PD-PD coupling) depends on, or is influenced by, their oscillation frequency and, if so, if the coupling also shows resonance. In the context of this manuscript, resonance is defined as a neuronal property that produces a maximum response to oscillatory input at a non-zero frequency. To compare any frequency dependence of the electrical coupling and that of the individual neurons, it was necessary to measure these two factors simultaneously. To do so, we arbitrarily designated the two PD neuron as pre- and postjunctional, injected a sweeping-frequency sinusoidal (ZAP) current into the prejunctional PD neuron and measured the voltage responses in both pre-and postjunctional PDs (Fig. 2). We then switched the pre and post designations and repeated the protocol. In the trials shown here, the ZAP function frequency is swept from 0.1 Hz to 4 Hz, a range that covers the natural burst frequency of PD neurons which is typically between 0.5 and 2.5 Hz. In several trials we also changed the direction of the frequency sweep to go from high to low frequency. There was no difference in our measurements when the direction of the sweeping frequency of the ZAP current was reversed.

**Figure 2.**
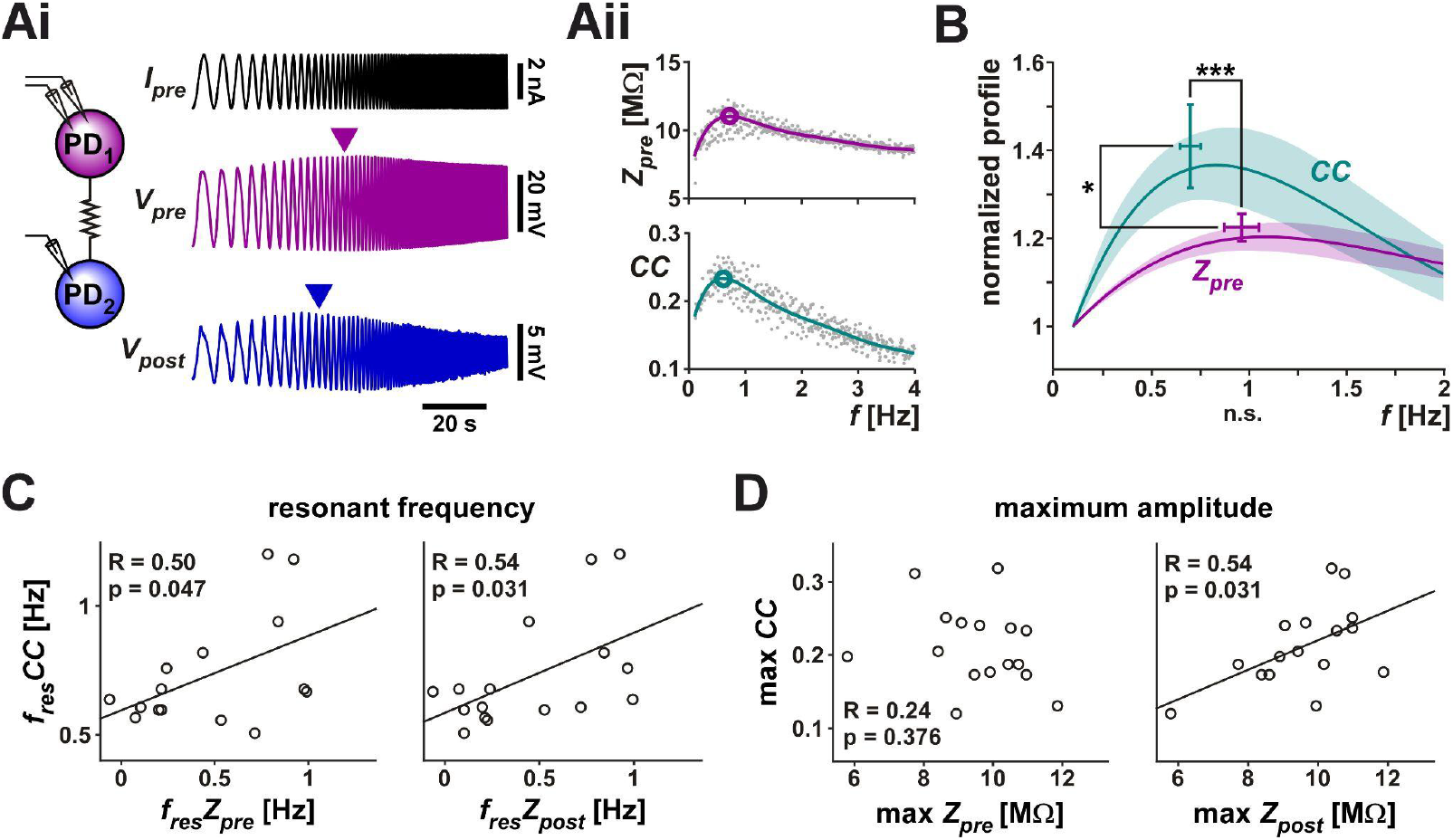
The coupling coefficient (*CC*) between the two PD neurons shows resonance. **(A)** A ZAP current, sweeping a frequency range of 0.1 to 4 Hz, was applied to one PD neuron to simultaneously measure the voltage changes in both PD neurons. **(Ai)** Both neurons showed a peak amplitude response at an intermediate frequency (marked by arrowheads). Schematic shows the two coupled neurons monitored in current clamp. **(Aii)** The prejunctional impedance (*Z*_*pre*_) and *CC* of the data shown in Ai. A 6^th^ order polynomial fit (smooth curves) to the raw data was used to measure the peak amplitude and resonance frequency (circled). **(B)** *Z*_*pre*_ and *CC* have distinct resonances. Averaged frequency profiles of *CC* and *Z*_*pre*_ are shown, both normalized to their amplitude at 0.1 Hz. *CC* had a smaller resonance frequency than *Z*_*pre*_ (p<0.001) and higher resonance power (p=0.037). N=19, paired Student’s t-test. **(C-D)** The resonance frequency of *CC* was correlated with the resonance frequency of both *Z*_*pre*_ and *Z*_*post*_ **(C)**, while its maximum amplitude was only correlated with that of *Z*_*post*_ **(D)**.

In 19 out of 28 measurements, both the prejunctional membrane impedance (*Z*_*pre*_; Table 1) and the coupling coefficient (*CC*) showed clear resonance (Fig. 2A). Note that the peak values shown in the figure do not exactly match the peak of the mean profile (solid line) since the peak of the average of multiple nonlinear curves is determined by the overall shapes of the individual curves, not just by their peaks. In response to the ZAP current, however, *Z*_*pre*_ and *CC* showed distinct frequency profiles (Fig. 2B): *CC* had a lower resonance frequency (0.70 ± 0.20 Hz) than *Z*_*pre*_ (0.97 ± 0.36 Hz) and the normalized peak amplitude of *CC* was larger than that of *Z*_*pre*_. Additionally, the resonance frequency of *CC* was correlated with the resonance frequency of both the prejunctional and postjunctional impedance (*Z*_*pre*_ and *Z*_*post*_, Fig. 2C), while its maximum amplitude was only correlated with that of *Z*_*post*_ (Fig. 2D).

### 3.1 Electrical coupling conductance shows a preferred frequency (resonance)

Membrane potential resonance can be measured using both current clamp and voltage clamp methods, each providing its own advantage. Current clamp measurements allow the membrane potential to change freely and therefore, voltage-dependent ionic currents can also influence the membrane potential. This method allows one to observe neuronal responses in a manner closer to their natural biological activity and, in general, current clamp measurements provide a more realistic value of the impedance amplitude (Rotstein and Nadim, 2019). However, because the electrical coupling coefficient is influenced by the input resistance of the postjunctional neuron, it is not a direct measure of the strength of electrical coupling (Bennett, 1966; Mann-Metzer and Yarom, 1999). A direct estimate of the electrical coupling conductance, *G*_*c*_, requires measuring the current flowing between the two coupled neurons (Table 1) and, to obtain an accurate measurement of the ionic current, the membrane potentials must be constrained using the voltage clamp method, as we showed in Figure 1C.

To directly measure whether the coupling conductance *G*_*c*_ is influenced by frequency, we voltage clamped both PD neurons at a holding potential of -60 mV. We then applied a ZAP function voltage waveform (ranging from -60 to -30 mV) to the prejunctional neuron, while holding the postjunctional neuron at a steady voltage of -60 mV (Fig. 3Ai). This allowed us to simultaneously measure the currents flowing in the pre- and postjunctional neurons (*I*_*pre*_ and *I*_*post*_) in response to the change in the frequency of *V*_*pre*_. As seen in the example in the figure, *I*_*pre*_ showed a clear minimum in response to the voltage ZAP, indicating a minimum in the neuronal admittance (the reciprocal of impedance) value. This simply reflects the membrane potential resonance in the prejunctional PD neuron as measured in voltage clamp (Tseng and Nadim, 2010)(Fig. 3Aii, top panel). Interestingly, in response to the prejunctional ZAP function, the postjunctional current, *I*_*post*_, did not remain constant in amplitude but had a clear maximum amplitude at a non-zero frequency. Therefore, the PD-PD coupling conductance, *G*_*c*_, also showed a peak at this frequency (Fig. 3Aii, bottom panel).

**Figure 3.**
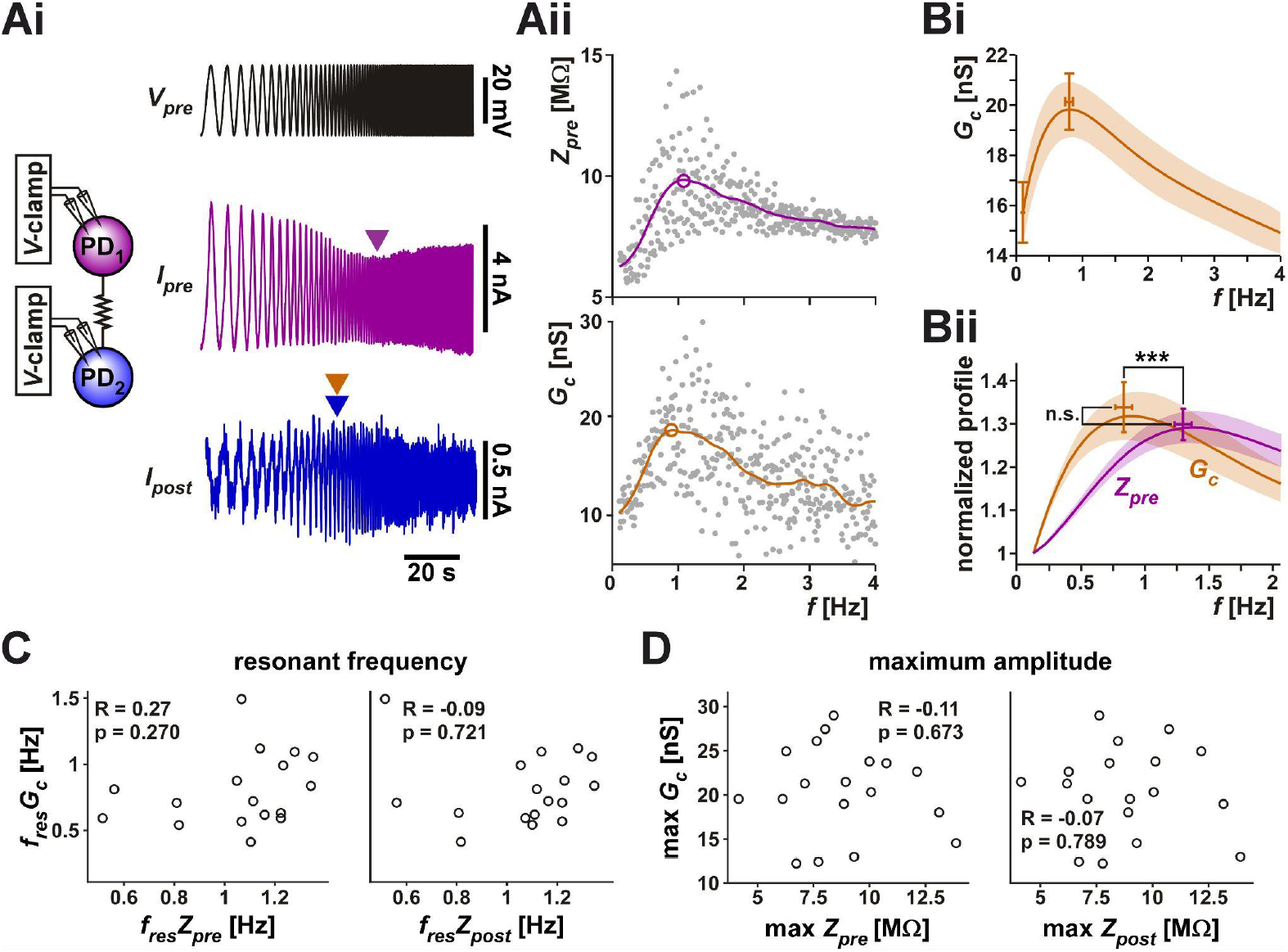
The coupling conductance shows a frequency-dependent resonance which is distinct from the resonance of the coupled PD neurons. **(A)** The two PD neurons were voltage clamped, the prejunctional neuron with a ZAP waveform, sweeping a frequency range of 0.1 to 4 Hz and a voltage range of -60 to -30 mV, while the postjunctional neuron was held at constant holding potential of -60 mV (not shown), and the current flow in both neurons was measured. (**Ai**) *I*_*pre*_ showed a minimum value at an intermediate frequency, reflecting the intrinsic resonance of the prejunctional neuron (magenta arrowhead), while *I*_*post*_ showed a peak at a distinct frequency (blue/bronze arrowheads). Schematic represents the two coupled neurons in voltage clamp. **(Aii)** The prejunctional impedance (*Z*_*pre*_) and *G*_*c*_ measured from the data shown in Ai. A 6^th^ order polynomial fit (smooth curves) to the raw data was used to measure the peak amplitude and resonance frequency (circled). The peak of *G*_*c*_ corresponds to the bronze color arrowhead in Ai. **(Bi)** The frequency profile of *G*_*c*_ across experiments shows a peak below 1 Hz. (**Bii**) *Z*_*pre*_ and *G*_*c*_ have distinct resonances. Averaged frequency profiles of *CC* and *Z*_*pre*_ are shown, both normalized to their amplitude at 0.1 Hz. *G*_*c*_ had a smaller resonance frequency than *Z*_*pre*_ (p<0.001) but comparable resonance power *Z*_*PD*_ (p=0.525). N=20, paired Student’s t-test. **(C-D)** Neither the resonance frequency **(C)**, nor the resonance amplitude **(D)** of *G*_*c*_ was correlated with that of *Z*_*pre*_ or *Z*_*post*_.

Unlike the measurements with the step protocol, in which the directionality of the electrical coupling had little influence, we found that the two directions of the coupling often produced slightly different results. Therefore, in this part of the study, we treated the PD1 to PD2 and the PD2 to PD1 in each preparation independently. In 20 of the 28 measured cases, *G*_*c*_ showed resonance. Fig. 3B shows the averaged resonance profile of these 20 electrical connections.

Because *Z*_*pre*_ and *G*_*c*_ have different units, their amplitudes cannot be directly compared. Yet it is useful to examine how much larger each of these factors is at its peak compared to its baseline. In fact, membrane potential resonance power is often measured as a ratio of the peak impedance *Z*_max_ to the impedance at zero frequency (i.e., the input resistance). We used the values of *Z*_*pre*_ and *G*_*c*_ at the lowest frequency (0.1 Hz) as a proxy for the zero-frequency values and normalized these curves to this value for each experiment (Fig. 3C). A paired comparison between *G*_*c*_ and the impedance profile (*Z*_*pre*_; see Table 1) of the prejunctional neuron showed no difference in their relative amplitudes. However, *G*_*c*_ had a significantly lower resonance frequency (0.80 ± 0.26 Hz) than *Z*_*pre*_ (1.27 ± 0.23 Hz). Also, note that the resonance frequencies for *Z*_*pre*_ were different between current clamp and voltage clamp experiments, because, as described above, *Z*_*pre*_ measured in current clamp is influenced by nonlinear actions of voltage-gated ionic currents. Finally, unlike with the coupling coefficient *CC*, we did not observe any correlation between *Z*_*pre*_ or *Z*_*post*_ and *G*_*c*_ either in frequency (Fig. 3D) or in amplitude (Fig. 3E). This is consistent with the hypothesis that *G*_*c*_ reflects the properties of the electrical coupling and not those of the coupled neurons.

### 3.2 Modeling elucidates how resonance of the coupling coefficient *CC* arises

Frequency dependence of electrical coupling may emerge from the properties of the coupled neurons, may be a property of the junctional coupling itself, or arise from the interaction of the two. To demonstrate how resonance of the coupling coefficient *CC* could arise from the membrane potential resonance properties of the coupled neurons, we coupled two biophysical models that capture the resonance properties of the isolated PD neuron (Fox et al., 2017) with a constant electrical coupling coefficient. We injected a ZAP current into one neuron and measured the voltage responses of both neurons (Fig. 4Ai). Current injection to PD model neuron 1 resulted in membrane potential resonance, mainly due to the intrinsic properties of this neuron, and current flow through the electrical coupling to PD model neuron 2 produces membrane potential resonance in the second neuron. In this simulation, the two PD model neurons were identical and therefore, when isolated, had identical impedance profiles (*Z*_1_ = *Z*_2_ in Fig. 4Aii). Coupling only slightly changed the impedance profile of prejunctional neuron 1 compared to its profile when isolated (*Z*_*pre*_ compared to *Z*_1_; see Table 1 for notations). In contrast, the impedance profile of the postjunctional neuron 2, when coupled, was quite distinct from its isolated profile (*Z*_*post*_ compared to *Z*_2_), because the current now flowed through the electrical coupling and was not directly injected into neuron 2. In this simulation, the *CC* vs. frequency curve also showed resonance, with a resonance peak frequency at a value very close to that of the coupled neurons. But what factors determined the resonance frequency of *CC*?

**Figure 4.**
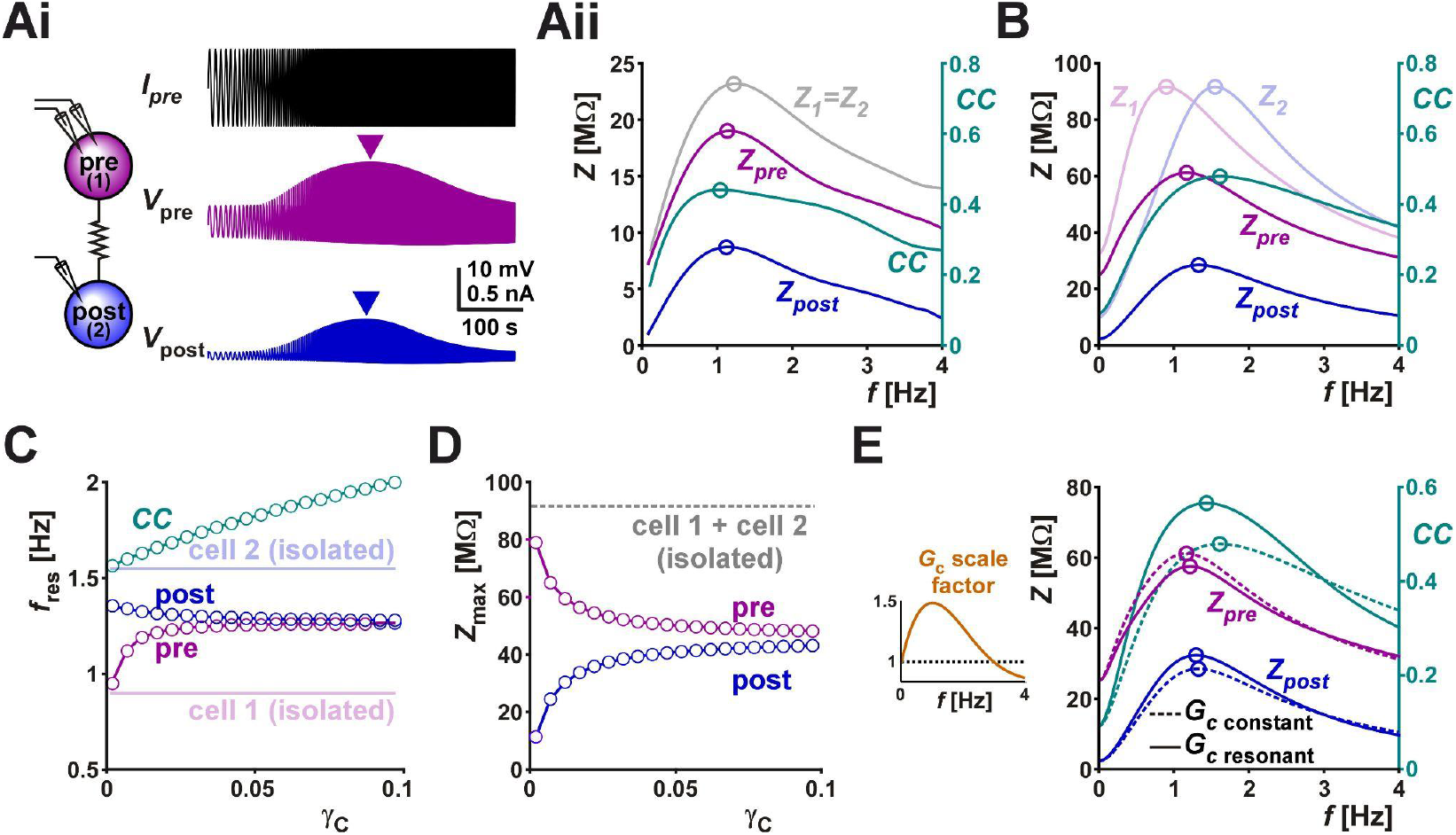
**(A)** Membrane impedance of the pre- and postjunctional PD model neurons (*Z*_*pre*_ and *Z*_*post*_, respectively) were measured by the response of the voltage amplitude to an oscillatory ZAP current input spanning 0.1 Hz to 4 Hz. Coupling coefficient (CC) was measured as the impedance profile of the pre (1) and post (2) synaptic cells are the same when synaptically isolated, shown as the gray line in Aii, and differ in amplitude when electrically coupled (shown as the purple line for *Z*_*pre*_ and the blue line in *Z*_*post*_). The coupling coefficient has a resonance frequency that is similar to the membrane impedance profiles. **(B)** (the analytical calculation) shown for when the isolated pre (1) and post (2) synaptic cells have different resonant frequencies, indicated as *Z*_1_ and *Z*_2_, and display an intermediate resonant frequency when electrically coupled. In contrast, the coupling coefficient resonant frequency does not take a value between the resonant frequencies of *Z*_1_ and *Z*_2_. **(C)** The resonant frequencies are shown as a function of increasing *γ*_*c*_, where the membrane impedance fRes converge to a value that is intermediate to the resonance frequencies of the isolated cells (indicated as cell 1 and 2). The coupling coefficient value increases monotonically as a function of *γ*_*c*_. **(D)** The maximal impedances for the isolated cells are equal (shown as dashed gray line, the same as in (B)), and approach a similar, lesser value as a function of increasing *γ*_*c*_. **(E)** The case of a frequency-dependent coupling conductance is considered, where it *G*_*c*_ is either at a fixed value (1, *G*_*c*_ constant, dashed line) or changes as a function of frequency (resonant, solid line). The membrane impedance profiles are compared in both cases, with a negligible effect on resonance frequency and amplitude for both *Z*_*pre*_ and *Z*_*post*_, with an effect of similar magnitude for the coupling coefficient.

To address this question, we switched to linear resonator neurons in which the impedance profile can be mathematically calculated (Richardson et al., 2003; Rotstein and Nadim, 2014). The full analysis is provided in Appendix 1. In the linear system of two coupled resonator neurons, the value of the coupled impedance profiles, as a function of the respective uncoupled profiles is given by

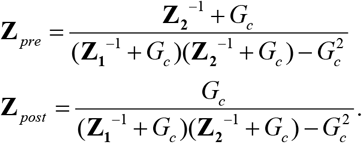

(equations (1.8) of the Appendix with notations of Table 1) and the value of *CC* reduces to the ratio of the amplitudes of the two impedance profiles.

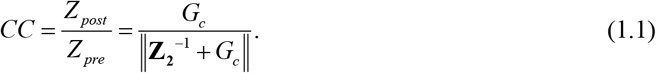

Here *f* is the input frequency and **Z**_2_ is the complex impedance profile of the postjunctional neuron when isolated (*Z*_2_ is the amplitude of the complex **Z**_2_, i.e., *Z*_2_ = ‖**Z**_2_‖). Note that for linear resonator neurons, *CC* only depends on the impedance of the (isolated) postjunctional neuron and not on that of the prejunctional neuron. Although this result does not generally hold for nonlinear (e.g., biological) resonators, it still provides a very good approximation in most cases in addition to a clearer conceptual understanding of the phenomenon.

The coupled linear resonators provide insight into how electrical coupling influences the resonance properties of the neurons as well as that of *CC*. For instance, coupling two linear resonators with the same maximal amplitude, but distinct resonance frequencies, shifted the resonance frequencies of both neurons toward values in between those of the isolated neurons (Fig. 4B; compare peak frequencies of *Z*_*pre*_ and *Z*_*post*_ with *Z*_1_ and *Z*_2_). The resonance frequency *Z*_*post*_ fell between *Z*_*pre*_ and *Z*_2_. The postjunctional impedance profile (*Z*_*post*_: which is *V*_2_/*I*_1_ in Figure Ai; see Table 1) always had a lower amplitude than the prejunctional profile (compare *Z*_*pre*_ and *Z*_*post*_ in Figure 4B). In Figure 4B, we also show the frequency-dependent profile of *CC* for comparison (note the different scales). Here, the resonance frequency of *CC* was close to that of *Z*_2_. Interestingly, however, the resonance frequency of *CC* was not constrained to fall between the resonance frequencies of *Z*_1_ and *Z*_2_. When the electrical coupling conductance was small, the resonance frequency of *CC* was close to that of *Z*_2_, but when *G*_*c*_ was increased, this frequency also increased monotonically (Fig. 4C). Not surprisingly, increasing the strength of coupling also caused the resonance frequencies (Fig. 4C) and maximum pre- and postjunctional impedance values (Fig. 4D) to converge to the same value.

We can also use the coupled linear resonators to predict how a frequency-dependent *G*_*c*_ may influence the measured coupling coefficient. To make this comparison, we scaled *G*_*c*_ as a function of frequency in a manner similar to what we had measured in the biological system (Fig. 3Bii and insert of Fig. 4E). A comparison of the resulting *CC* and the *CC* obtained with a constant *G*_*c*_ value across frequencies showed that frequency dependence of *G*_*c*_ can clearly amplify the amplitude of *CC* near the resonance frequency of *G*_*c*_, by bringing the *Z*_*pre*_ and *Z*_*post*_ curves closer to each other in this range (Fig. 4E).

### 3.3 Can coupling conductance resonance result from network connectivity?

When both neurons are voltage-clamped, the prejunctional neuron with a fixed-amplitude sinusoidal waveform and the postjunctional neuron at a constant holding voltage, the amplitude of the ionic current change recorded in the postjunctional neuron (*I*_*post*_ or the coupling current) is proportional to the coupling conductance *G*_*c*_ and independent of any resonant properties of either neuron. This follows from the fact that where *V*_*pre*_ and *V*_*post*_ are controlled by voltage clamp and *G*_*c*_ is constant. Although this is an obvious result, it is informative. We demonstrated this in the simulation shown in Figure 5A, where the prejunctional model neuron was voltage-clamped with a ZAP function (range -60 to -45 mV) and the prejunctional neuron was held at a steady voltage of -60 mV. The current (*I*_*pre*_) in the prejunctional neuron showed a minimum, while the current flowing to the postjunctional neuron (*I*_*post*_) did not change with the frequency of the ZAP function. This is also clear from our calculations for the coupled linear resonators in voltage clamp as shown in Appendix 1 (see equation (1.11)).

**Figure 5.**
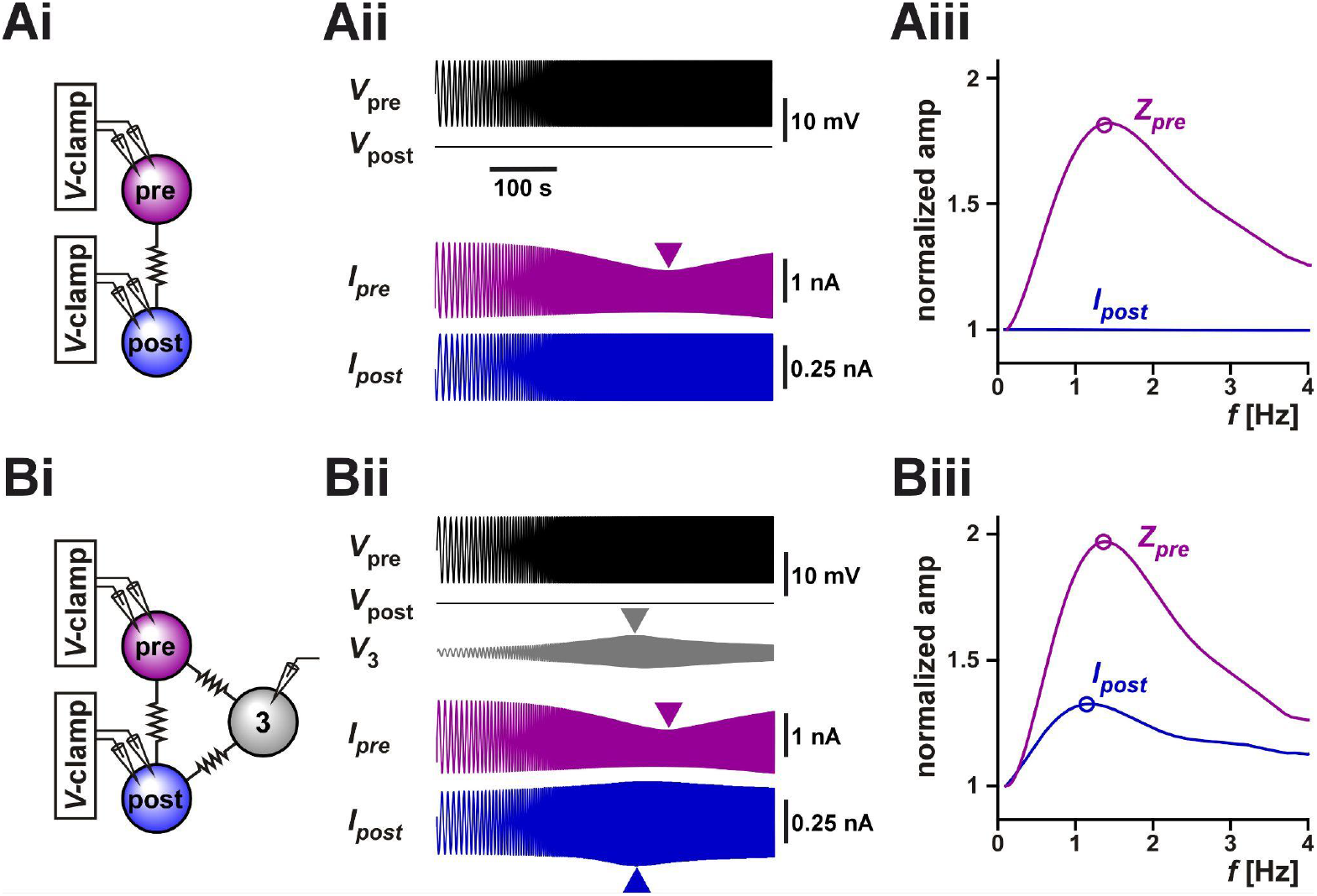
Coupling to a third resonant neuron can produce resonance in the coupling current between two voltage-clamped neurons. **(A)** The coupling current between two identical model neurons with resonant properties was measured in voltage clamp (schematic in **Ai**). The prejunctional neuron was voltage clamped with a ZAP waveform spanning from 0.1 Hz to 4 Hz and voltage range of -60 to -45 mV. The postjunctional neuron was voltage clamped at a holding potential of -60 mV. The postjunctional current amplitude showed no frequency dependence **(Aii)**. As a function of input frequency, the prejunctional impedance shows resonance, but the post junctional current remains constant. For comparison, *Z*_*pre*_ and *I*_*post*_ are normalized to their value at 0.1 Hz. **(B)** The same protocol as A, but the two neurons are both coupled to a third (identical) neuron which is not voltage clamped (schematic in **Bi**). The addition of the third cell leads to a frequency-dependent response in the voltage of the third neuron **(Bii)** and in resonance in the postjunctional current **(Biii)**. For comparison, *Z*_*pre*_ and *I*_*post*_ are normalized to their value at 0.1 Hz.

However, when these two neurons are part of a circuit of electrically coupled neurons, even when both neurons are voltage-clamped, the measurement of *I*_*post*_ may not have a constant amplitude at all input frequencies due to circuit connectivity. For example, if both neurons are electrically coupled to a third neuron whose voltage can vary freely, indirect current flow through the third neuron may affect the amplitude of *I*_*post*_. Indeed, in the pyloric circuit, the two biological PD neurons are electrically coupled to the anterior burster (AB) neuron (Marder and Eisen, 1984) and, in our experiments described above, we did not control or monitor the activity of the AB neuron. It is therefore possible that the apparent resonance we observed in our experimental measurement of *I*_*post*_ (Fig. 3Ai) was due to the uncontrolled changes in the voltage of the AB neuron. To test this possibility, we coupled the model neurons of Figure 5A to a third neuron with the same resonance properties and ran the same voltage clamp protocol. Indeed, we observed that even though the pre- and postjunctional neurons were voltage-clamped, the voltage of the third coupled neuron (marked 3 in Fig. 5B) showed a peak at an intermediate frequency. Thus, the resonance of neuron 3 resulted in an apparent resonance in our measured *I*_*post*_, because in this case

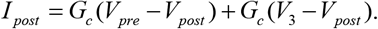

A normalized comparison between the impedance profile *V*_*pre*_ and *I*_*post*_ (Fig. 5Biii) shows that even when the three neurons are identical in their properties (and therefore have the same isolated resonance frequency), *I*_*post*_ may show resonance at a different frequency, as we had observed in our experimental measurements of Fig. 3A-B. Therefore, a potential mechanism for electrical coupling current resonance is through frequency preference inherited from other electrically coupled cells.

### 3.4 Potential function of electrical coupling resonance

We used computational modeling to understand the potential function of resonance in the electrical coupling conductance in this system. We used a computational model of two electrically coupled bursting neurons and chose the parameters of the two neurons to produce bursting oscillations with different cycle frequencies when uncoupled. We then coupled the two neurons and examined the synchronization of their activity at different electrical conductance strengths. The level of synchronization was measured as the coefficient of determination (R^2^) between the two voltage waveforms (Lane et al., 2016). We measured the synchrony of the full bursting waveforms between the two neurons (full). In addition, we lowpass-filtered the traces to measure the synchrony of only the slow waves (slow), and high pass-filtered to measure the synchrony of only the spiking activity (fast).

To examine the effect of resonance in *G*_*c*_ on the synchrony between the two neurons, we produced a *G*_*c*_ frequency profile similar to that observed experimentally (Fig. 6A; compare with Fig. 3Bii). Although the two model neurons had different intrinsic burst frequencies, they always oscillated with the same frequency (i.e., they were phase locked) when coupled. To understand the role of *G*_*c*_ resonance, we changed this burst frequency by modifying the intrinsic properties of the bursting neurons (see Methods). We found that when the two cells oscillated at either low or high frequencies, where the *G*_*c*_ was smaller, the slow wave synchrony between the two neurons was smaller (Fig. 6Bi, Biii, and C). In contrast, when the network frequency matched the *G*_*c*_ resonance frequency, the level of synchronization was maximal (Fig. 6Bii and C). In contrast to the slow wave, the fast spiking activity of the two neurons was not noticeably altered by frequency. When *G*_*c*_ was kept constant as a function of frequency, then network frequency did not affect the level of synchrony between the two neurons, either in the slow wave or in the spiking activity. The level of synchrony in this case was determined simply by the value of the electrical coupling conductance *G*_*c*_ (Fig. 6D).

**Figure 6.**
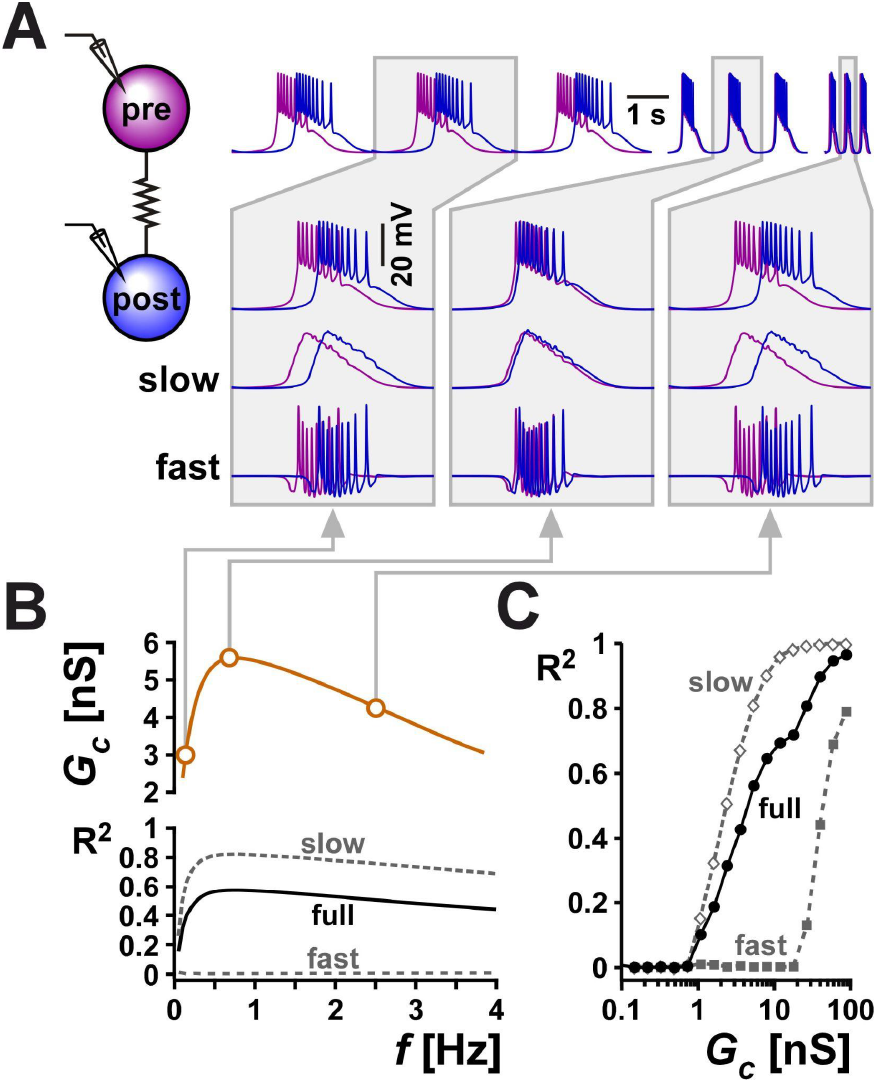
Resonance in the coupling conductance influences the level of synchrony between two model bursting neurons. (**A)** The level of synchrony between two model bursting neurons, coupled with a resonant *G*_*c*_ (schematic), depends on the network oscillation frequency. The three columns show superimposed phase-locked oscillations of two model bursting neurons at three frequencies. The second row is a zoom in to a single burst. The third row shows lowpass filtered traces (slow), highlighting the level of asynchrony of the burst slow waves. The bottom row shows the high pass filtered traces (fast = full - slow), highlighting the lack of synchrony of spiking activity. Gray boxes correspond to frequencies and *G*_*c*_ values as shown in panel B. **(B)** Coupling conductance is modeled to show resonance at *f* = 0.75 Hz. The level of synchrony between the two coupled neurons, measured as a coefficient of determination R^2^ of their voltage waveforms depends on the network frequency. Changing the network frequency increases synchrony of the slow and full waveforms, but not the fast spiking activity. (**C)** R^2^ increases with the coupling conductance. Tables`

## 4 Discussion

Gap junction-mediated electrical coupling between neurons is well known to lead to synchronization of their electrical activity (Gutierrez et al., 2013; Marder et al., 2017; Alcamí and Pereda, 2019; Vaughn and Haas, 2022). However, as a number of modeling studies have shown, in certain conditions it can also promote anti-synchrony (Sherman and Rinzel, 1992; Chow and Kopell, 2000; Bem and Rinzel, 2004). It is commonly assumed that electrical coupling acts primarily as a lowpass filter so that slow voltage changes, such as burst envelopes and subthreshold oscillations, are transmitted more effectively than fast ones such as action potentials (Galaretta and Hestrin, 1998; Connors and Long, 2004; Placantonakis et al., 2006). However, more recent studies that have explored electrical coupling in oscillatory networks have found that the interaction the intrinsic properties of neurons and the electrical coupling could result in a band-pass filtering of the coupling coefficient, such that the coupling coefficient is highest around a “resonance” frequency (Armstrong-Gold and Rieke, 2003; Curti et al., 2012; Stagkourakis et al., 2018). Such bandpass-filtering has been attributed to the properties of voltage-gated ion channels or subthreshold resonance in the coupled neurons (Curti et al., 2012; Alcamí and Pereda, 2019), thus suggesting that the subthreshold resonance frequency can play a significant role in setting the frequency of a network of electrically coupled neurons.

Here, we found similar results in the PD neurons of the crab pyloric circuit. The two PD neurons produce ongoing synchronous bursting activity, are strongly electrically coupled (Fig. 1) and show membrane potential resonance (Figure 2 and Tohidi and Nadim, 2009). We found that the coupling coefficient of these neurons also shows resonance, but at a much lower frequency than that of their membrane potential resonance (Fig. 2). The *CC* resonance frequency, however, was strongly correlated with both that of the pre- and postjunctional neuron. A combined modeling and mathematical analysis showed that although with increased coupling strength the resonance frequencies measured in the coupled neurons converges to the same value, the *CC* resonance frequency does not necessarily fall between these two values (Fig. 4C). In fact, our mathematical calculations, based on coupled linear resonators, showed that in response to oscillatory input, *CC* behaves very much like it does in response to a direct current input: It depends on a nonlinear combination of the coupling conductance and the impedance of the postjunctional, but not prejunctional, neuron (Equation (1.1); also see (Alcamí and Pereda, 2019)). Thus, at least to the first order (linear) approximation, the resonance properties of the prejunctional neuron have no influence on the *CC* resonance frequency, which can fall well outside the range of resonance frequencies of the neurons. This finding is important in the light of the above-mentioned fact that *CC* resonance frequency is often considered to be a determinant of the network oscillation frequency (Curti et al., 2012; Stagkourakis et al., 2018).

The second, perhaps more surprising, finding of our study is that when we measured the current flow between the coupled PD neurons in voltage clamp, we found that the measured coupling was both frequency-dependent in its amplitude and had a resonance frequency distinct from the intrinsic resonance of the PD neurons. For direct current flow between voltage-clamped coupled neurons, this finding inevitably leads to the conclusion that the coupling conductance *G*_*c*_ is frequency-dependent. There are some caveats, however, that should be considered when drawing such a conclusion. First, voltage clamp is often subject to lack of space clamp. If gap junctions that lead to electrical coupling are present in a distal location from the voltage-clamped somata, it is possible that space clamp issues may somehow result in the appearance of frequency-dependence in the coupling current. Although we did not show these results in this manuscript, a structured multi-compartmental model of the coupled neurons did not show any significant resonance in the measured coupling current. This is consistent with previous findings showing that the stomatogastric neurons are quite electrotonically compact (Otopalik et al., 2019). The second caveat in drawing a conclusion that *G*_*c*_ is frequency-dependent is that both PD neurons are strongly coupled to the pyloric pacemaker AB neuron, which was neither voltage-clamped nor photo ablated (Miller and Selverston, 1979) here. In fact, a computational model of the three-neuron coupled circuit showed that a free-running AB neuron may indeed result in an apparent resonance of the coupling current measured between the two PD neurons (Fig. 5B). Although we did not resolve the caveat of coupling to additional neurons in the current study, our unpublished results indicate that there is a possibility that frequency-dependence may in fact in part be inherent to the electrical coupling conductance. These findings showed that peptide neuromodulators that activate the same ionic current in the pyloric pacemaker neurons have an opposite effect on shifting both the frequency and amplitude of resonance in the coupling current (Li et al., 2017). This result cannot be explained by coupling to a free-running AB neuron which is modulated the same way by the two peptides. Consequently, the gap junction channels may in fact have kinetics that allows for bandpass filtering. Although it is know that current flow through gap junctions may have complex and functional voltage-dependent properties (examples in Coleman et al., 1995; Vaughn and Haas, 2022), to our knowledge, such a frequency-dependent filtering property of gap junctions has not been previously reported.

Previous studies have suggested that different resonant properties of different circuit components collectively influence network frequency (Lovett-Barron et al., 2017). However, it remains to be determined to what extent *CC* or *G*_*c*_ resonance interacts with other frequency-dependent properties of a network. We showed, however, that resonance in *G*_*c*_ or the coupling current would amplify the resonance properties of *CC* (Fig. 4E). In addition, one functional consequence of the frequency-dependence of the coupling is intuitively clear if the network frequency may be subject to context-dependent changes. We demonstrated this using a coupled network of two intrinsically distinct model neurons. Although at all frequencies tested, the two neurons remained phase locked, their degree of synchronization was effectively determined by the frequency-dependent properties of the coupling conductance (Fig. 6). In an oscillatory network such as the crab pyloric network, where network frequency depends on multiple factors including neuromodulation and temperature, it is reasonable to assume that the degree of synchronization between the PD neurons may be influenced indirectly by the factors that modify network frequency. Although the experimental verification of these functional consequences remains to be performed, our combined experimental and modeling findings indicate that the resonance properties of electrical coupling may play a central role in shaping the output of oscillatory networks.

## 6 Acknowledgments

The authors thank Dr. Jorge Golowasch for his input and comments.

## 7 Appendix 1

### 7.0 Electrically coupled linear cells receiving oscillatory inputs

The general form of the electrically coupled two-cells network model we use is given by

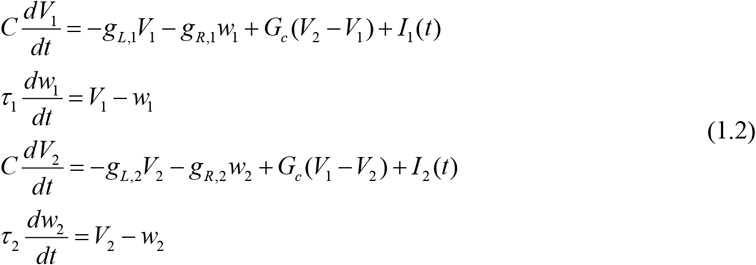

The dynamics of the individual cells in system (1.2) are the linearization of biophysically plausible (conductance-based) models around the resting potentials (Richardson et al., 2003; Rotstein and Nadim, 2014; 2019). For *k* = 1,2, *V*_*k*_ represents the membrane potential for the two cells and measures deflections from a resting potential (which here would be equal to 0), *w*_*k*_ represents the corresponding recovery variables after linearization, *t* (ms) is time, *C* is the specific capacitance, *g*_*L,k*_ are the linearized leak conductances, *g*_*R,k*_ are the linearized ionic conductances, *G*_*c*_ (µS/cm^2^) is the electrical coupling conductance, and *I*_*k*_ are time-dependent currents. In this Appendix, we are using dimensional parameters, with time in ms, frequencies in Hz, voltages and recovery variables in mV, capacitance in µF/cm^2^, conductances in µS/cm^2^ and currents in mA/cm^2^.

In current-clamp (I-clamp),

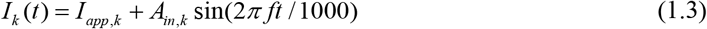

for *k* = 1, 2, where *A*_*in,k*_ and *f* are the externally-applied amplitudes and frequencies and *I*_*app,k*_ is a constant (DC) current. In voltage-clamp (V-clamp),

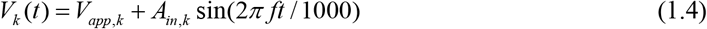

for *k* = 1, 2, where *A*_*in,k*_ and *f* are as above and *V*_*app,k*_ is a constant holding voltage. In the cases we consider here, except for the uncoupled cells (*G*_*c*_ = 0) that we use as a reference case to establish the resonant properties of the individual cells, only one cell (cell 1) receives an oscillatory input (regardless of whether it is in I- or V-clamp). Therefore, we refer to cells 1 and 2 as the pre- and postjunctional cells, respectively. To simplify the notation, we define

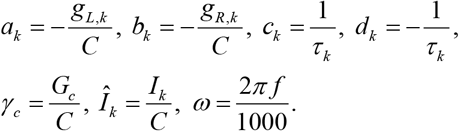

Substitution into system (1.2) yields

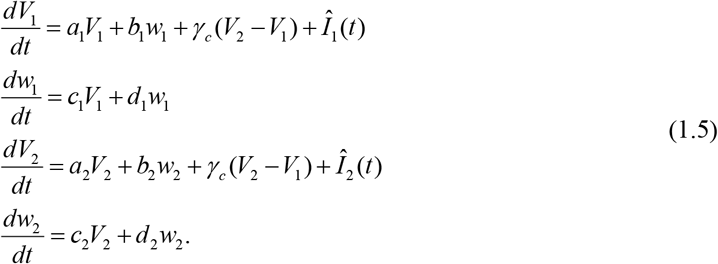

For use below, we further define the determinants and traces of the matrices (for *k* = 1, 2) of the coefficients of the linear system:

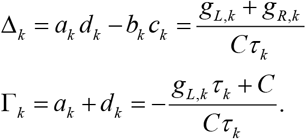

### 7.1 Response of the uncoupled cells to oscillatory inputs: cellular impedances and inverse admittances

Here we consider *γ*_*c*_ = 0 and *Î*_*k*_ (*t*) given by (1.3), with *A*_*in*,1_ = *A*_*in*,2_ = *A*_*in*_ and *I*_*app*,1_ = *I*_*app*,2_ = 0. The impedances of the individual uncoupled cells, as described previously (Richardson et al., 2003; Rotstein and Nadim, 2014), are given by

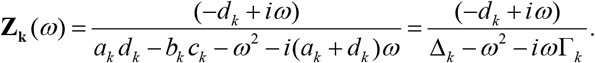

The impedance amplitudes and phases (phase-shifts) are given, respectively, by

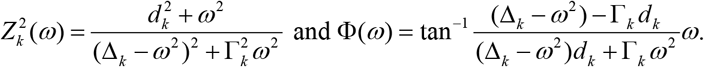

Therefore, the solutions to equations (1.5) for the uncoupled neurons, each receiving sinusoidal input currents, are given by

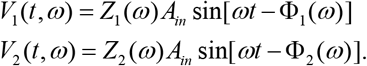

These calculations correspond to I-clamp. In V-clamp, *V*_*k*_(*t*) is given by (1.4) *A*_*in*,1_ = *A*_*in*,2_ = *A*_*in*_ and *V*_*app*,1_ = *V*_*app*,2_ = 0. Since the system is linear, as described previously (Rotstein and Nadim, 2019), the admittances are given by

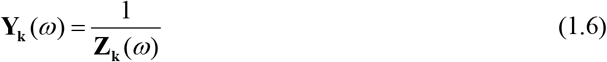

and

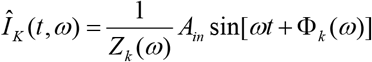

for *k* = 1, 2. Note that for nonlinear systems, the equality between the impedance (measured in I-clamp) and the inverse admittance (measured in V-clamp) does not generally hold Rotstein, 2019 #4352}.

In order to compute the impedances, we used the complex exponential expression for *Î*_*k*_ (*t*) = *A*_*in*_exp(*iωt*) and assumed (from linearity) that the stationary solutions to system (1.5) are given by

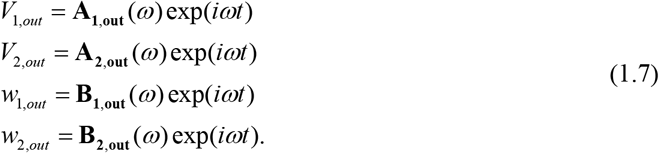

We then substituted these expressions into equations (1.5) and computed the coefficients

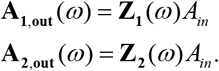

### 7.2 Response of the electrically coupled cells to oscillatory inputs solely to the prejunctional cell (cell 1) in I-clamp

Here we assume that *Î*_1_ (*t*) is a sinusoidal input current of the form (1.3) with *A*_*in*,1_ = *A*_*in*_, *I*_*app*,1_ = 0 and *Î*_2_ (*t*) = 0. Equivalently, *Î*_1_ (*t*) = *A*_*in*_exp(*iωt*) and *Î*_2_ (*t*) = 0. Substitution of the formal solutions (1.7) into equations (1.5) yields

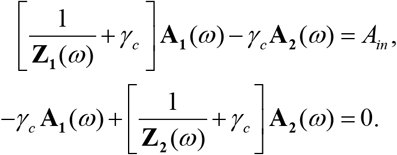

By solving this algebraic system, we obtain

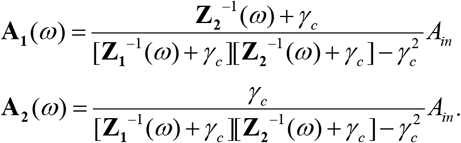

Therefore, the impedances of the coupled cells are given by

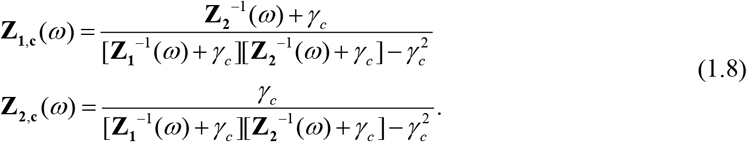

The corresponding solutions to system (1.5) are given by

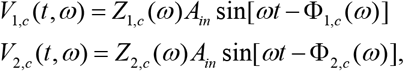

where *Z*_*k,c*_(*ω*) and Φ_*k,c*_(*ω*) are the amplitudes and phases of **Z**_*k,c*_(*ω*) for *k* =1, 2. We refer to *Z*_1,*c*_ as the prejunctional impedance and to *Z*_2,*c*_ as the postjunctional impedance (*Z*_*pre*_ and *Z*_*post*_ respectively in Table 1).

These calculations assume the postjunctional cell (cell 2) is I-clamped. If, instead, the postjunctional cell is V-clamped, (*V*_2_(*t*) = *V*_*app*,2_), then

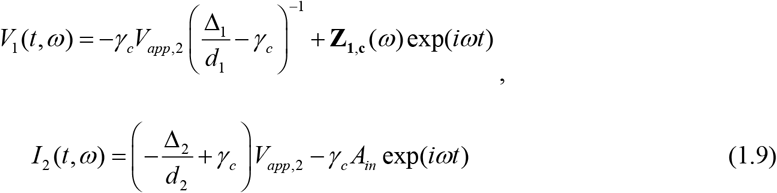

with

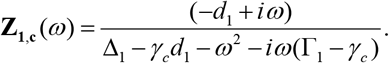

Therefore

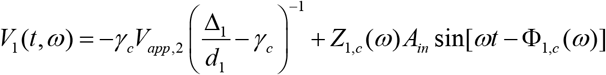

with

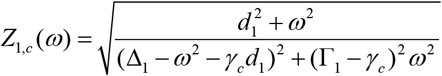

and

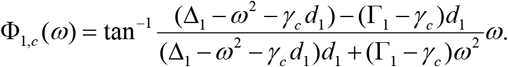

From (1.9),

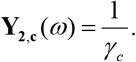

### 7.3 The coupling coefficient, *CC*

The coupling coefficient (*CC*; Table 1) is given by

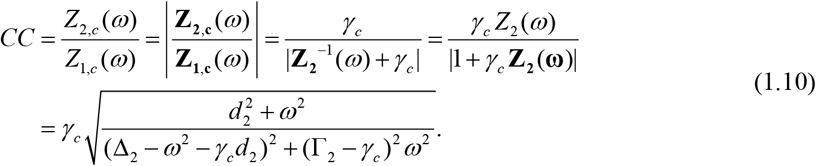

Formally, *CC* can be expressed in terms of the impedance of the isolated postjunctional cell and is independent of the impedance of the prejunctional cell.

If *Z*_2_(*ω*) acts as a low-pass filter (i.e., *b*_2_ = 0), then

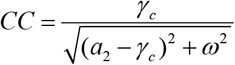

is also a low-pass filter.

### 8 Response of the electrically coupled cells to oscillatory inputs solely to the prejunctional cell (cell 1) in V-clamp

Here we assume that *V*_1_(*t*) is a sinusoidal input of the form (1.4) with *A*_*in*,1_ = *A*_*in*_ and *V*_*app*,1_ = 0 and *V*_2_ = *V*_*app*,2_ at a constant value. Equivalently, *V*_1_(*t*) = *A*_*in*_ exp(*iωt*). Substitution of these expressions into (1.5) yields

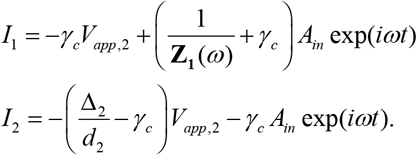

Therefore, the admittance (1.6) of the coupled neurons are given by

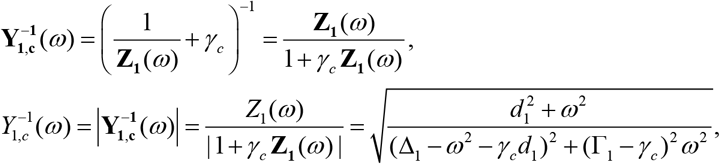

and

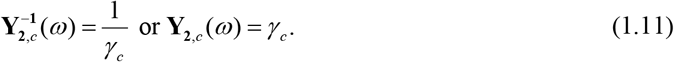

## 9 Data Availability Statement

### 10 Author Contributions

XL and FN conceived and designed the experiments and analysis. XL, performed all experimental analysis. OI, HR and FN designed and performed the computational modeling. HR performed all mathematical analysis. XL and FN wrote the manuscript draft. All authors contributed to the conceptual understanding of the findings and edited the manuscript.

### 13 Supplementary Material

None.

